# Molecular profiles and mutation burden analysis in Chinese patients with gastric carcinoma

**DOI:** 10.1101/449736

**Authors:** Chao Chen, Chunmei Shi, Xiaochun Huang, Jianwei Zheng, Zhongyi Zhu, Qiaolian Li, Si Qiu, Zhiqing Huang, Zhenkun Zhuang, Riping Wu, Panhong Liu, Fan Wu, Shanyun Lin, Bo Li, Xiuqing Zhang, Qiang Chen

## Abstract

The goal of this work was to investigate the molecular profiles and mutation burden in Chinese patients with gastric carcinoma (GC). In total, we performed whole exome sequencing (WES) on 74 GC patients with tumor and adjacent normal formalin-fixed, paraffin-embedded (FFPE) tissue samples. The mutation spectrum of these samples showed a high concordance with TCGA and other studies on GC. We found the alterations of 17 DNA repair genes (including BRCA2, POLE and MSH3, etc.) were strongly correlated with the tumor mutation burden (TMB) and tumor neoantigen burden (TNB) of GC patients. Patients with mutations of these genes tend to have high TMB (median of TMB = 12.77, p=2.3e-6) and TNB (median of TNB = 5.97, p= 2.8e-3). In addition, younger GC patients (age < 60) have lower TMB (p = 0.0021) and TNB (p = 0.034) than older patients (age >= 60). Furthermore, we found a list of 18 genes and two genomic regions (1p36.21 and Xq26.3) were associated with peritoneal metastasis (PM) of GC, and patients with amplification of 1p36.21 and Xq26.3 have a worse prognosis (p=0.002, 0.01, respectively). Our analysis provides GC patients with potential markers for single and combination therapies.

---

Gastric carcinoma (GC) is one of the most common cancers and a leading cause of cancer death worldwide ^1^, with a 5-year survival rate of about 30% ^2^. The highest incidence is in East Asia, Central and Eastern Europe, and South Africa ^3^. Surgery, chemotherapy and radiotherapy are the mainstay treatments of GC, but nearly 20% GC patients develop peritoneal metastasis (PM), which has a poor prognosis ^4^.

Several studies have used next generation sequencing strategies to determine the mutation spectrum of GC, and many significantly mutated driver genes have been identified, such as *TP53, ARID1A, PIK3CA*, and others ^5-8^. GC is divided into several subtypes according to its molecular classification, such as the MSI-high types which contains hypermutated samples and can be used as a potential marker for checkpoint-blockade therapy ^7^. In addition to MSI status, there are several other factors considered to be associated with responses to immunotherapy, such as PD-1/PD-L1 expression, mismatch repair deficiency, TMB and TNB ^9^. Rosenberg and colleagues reported that TMB was more significantly related to the response rates than the expression of PD-L1, suggesting the application prospect of TMB in cancer immunotherapy ^10^. However, most of these studies have been performed using fresh frozen (FF) tissues, but FF tissue has limited availability; therefore, our knowledge of GC and its treatment are far from complete ^11^. Formalin-fixing paraffin-embedding (FFPE) has been a standard sample preparation method for decades, and they are useful resources for cancer studies. There are many efforts to develop strategies to use FFPE specimens in cancer research, and several studies confirmed the technical feasibility ^12-14^. However, these studies mainly use next-generation sequencing (NGS) target region panels, and whole exome sequencing (WES) has rarely been reported in studies with a large sample size.

In this study, we first performed WES on 74 FFPE samples of GC based on the BGISEQ-500 platform, compared the molecular profiles of Chinese southern GC patients with TCGA and other cohorts, and then investigated the TMB and TNB of them. We found a panel of 17 DNA repair genes associated with high TMB and TNB, which can be used as potential markers for immunotherapy. Last but not least, we also discovered 18 genes and 2 regions related with the PM of GC, which can be further validated in large-scale studies.

## Results

### Patient characteristics

A total of 74 paired normal and tumor samples were successfully sequenced; 28 (38%) were less than 60 years of age, 46 (62%) were more than 60 years of age. The majority of the subjects were male (52,70%), and the remaining 22 (30%) were female. In all 7 (9%) were stage I, 8 (11%) stage II, 51 (68%) stage III, and 9 (12%) stage IV; and 26 (35%) patients had peritoneal metastasis in a follow-up exam. The clinical characteristics and statistics were list in **Table 1** and **Supplementary Table S1.**

**Table 1.**
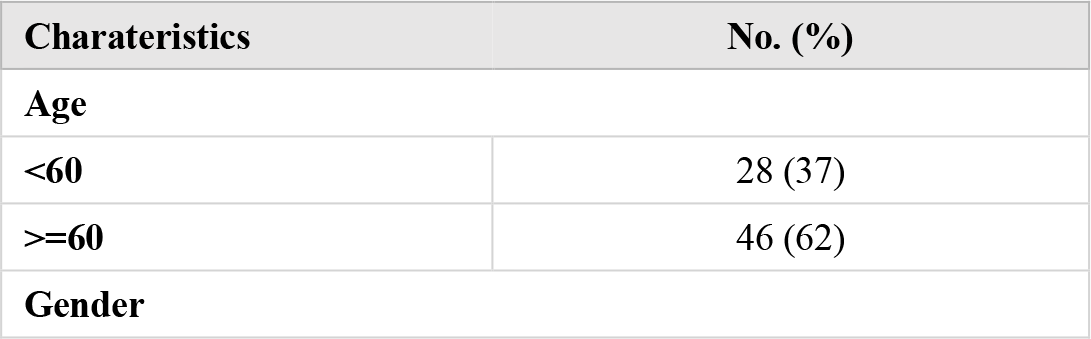
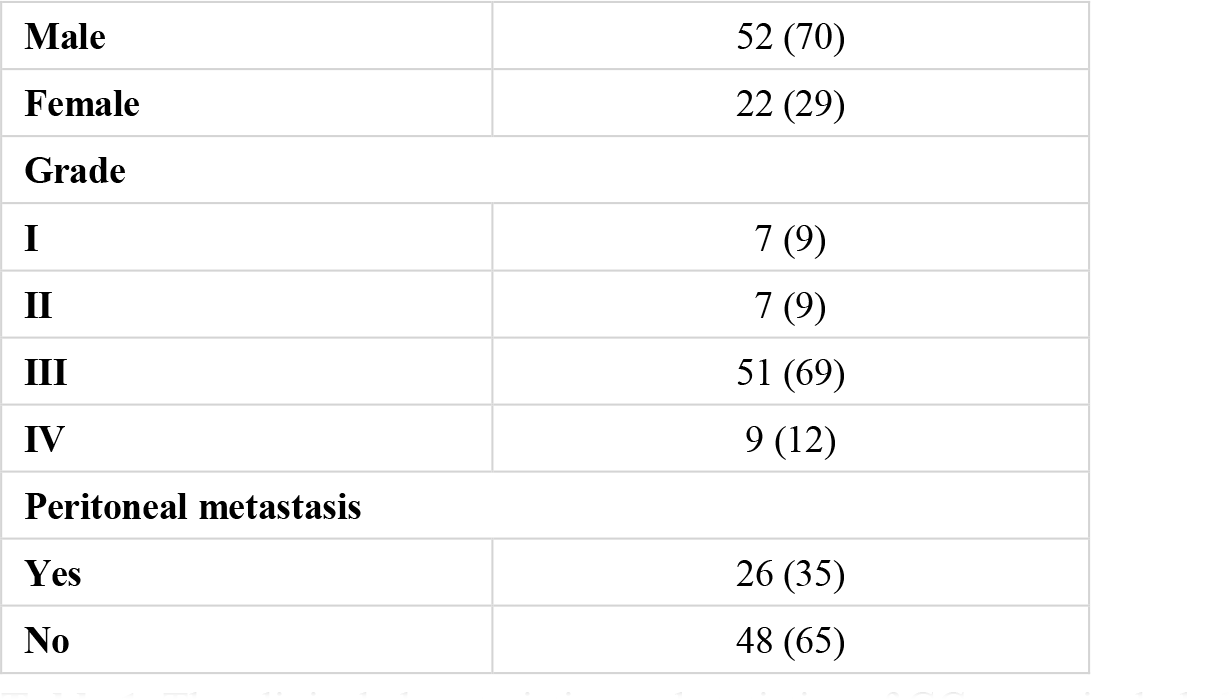
The clinical charateristics and statistics of GC cases included in this study (N = 74)

### Genomic profiles of Chinese GC patients

A total of 11,118 mutations were detected in this study, the mean number of somatic mutations per patient was 150 (range from 0 to 1517) (**Supplementary Table S2).** Somatic SNVs (sSNVs) and indels (sIndels) accounted for 95.4% and 4.6% of the mutations, respectively. Of the mutations, 3,066 (27.6%) were synonymous, 6,857 (61.7%) missense, 463 (4.2%) nonsense (stopgain), 9 (0.1%) stoploss, 212 (1.9%) splice site, 452 (4.1%) were frameshift indels, and 59 (0.5%) were in-frame indels. Several cancer-related genes were frequently mutated in our cohorts, such as *TP53* and *ARID1A*, consistent with previous studies on GC ^7,8,15^ (**Fig. 1A, Supplementary Table S3**). We randomly selected 36 mutation sites for mass spectrometry validation, and 34 (94.4%) of them were verified as somatic mutations (**Supplementary Table S4**).

**Figure 1:**
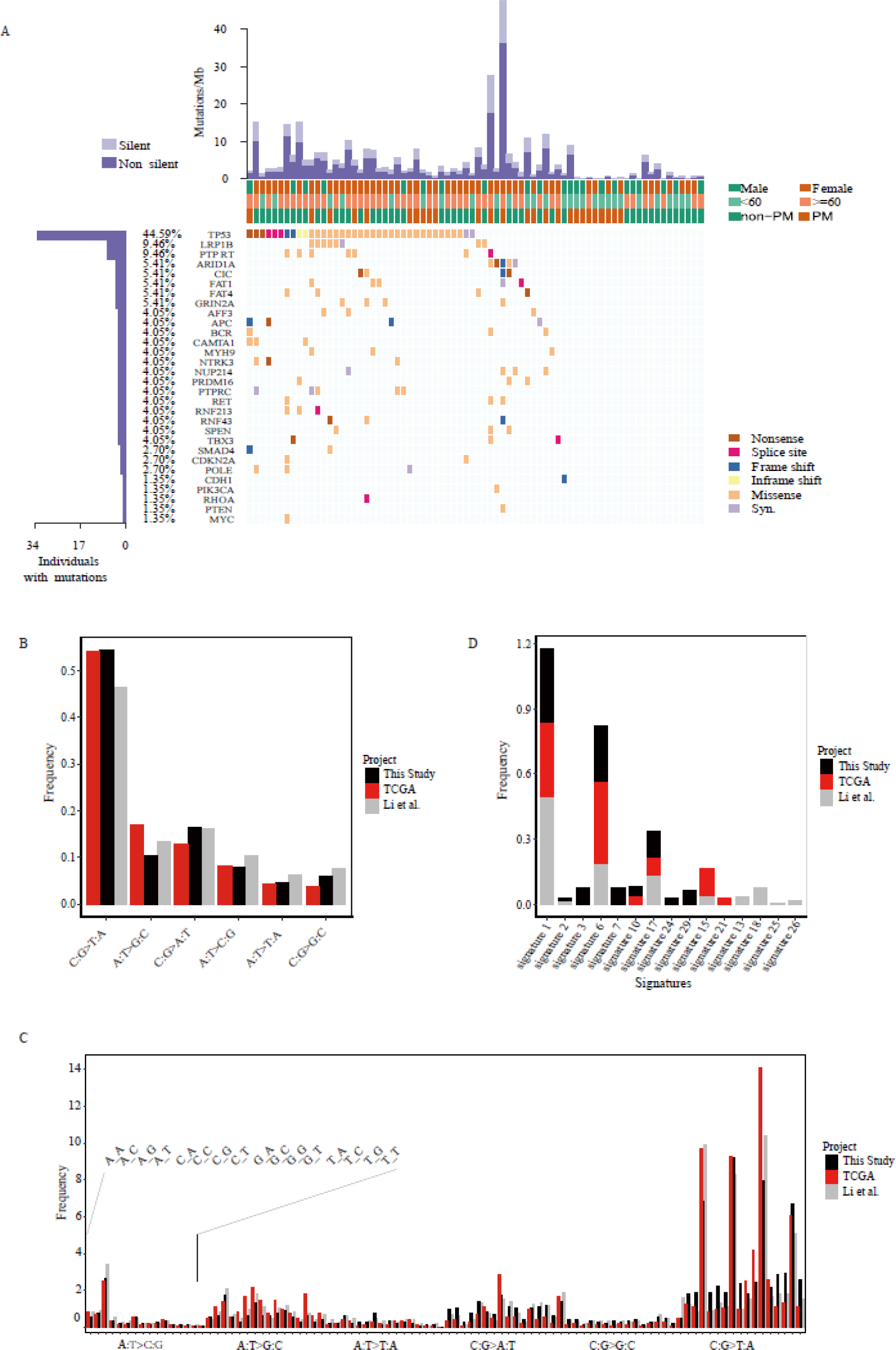
The mutation spectrum of GC in this study and the comparation with other studies. **(A)** Heat map showing somatic mutation profiles of cancer genes in this study. Left, the number of samples with mutations in a given gene. Top, the clinical type of samples and mutations burden of each sample. **(B)** The six classes of base substitution in three studies. **(C)** The 96 context-dependent (5’ to 3’) mutations patterns in three studies. **(D)** Each signature is displayed according to the 96 context-dependent mutation patterns in three studies.

The mutations in the exon and splice site regions of other two datasets, TCGA (download from https://cancergenome.nih.gov/) and Li et al. ^15^ were used for further comparative analysis. For point mutations, C > T, G > A transitions accounted for 54.4 % of the sSNVs, and the ratio of the 6 types of base substitution is similar to the studies of TCGA and Li et al. (**Fig. 1B**). We further found that the spectrum of flanking nucleotides surrounding the mutated base was highly concordant between our results and the other two datasets (**Fig. 1C**). The context-dependent mutational patterns of these three datasets were then identified using mSignatureDB (http://tardis.cgu.edu.tw/msignaturedb/) to explore the heterogeneity of mutagenic processes in GC and its diagnostic potential ^16^. The results showed that prevalence of signatures 1, 6, and 17 were similar in the three studies, accounting for the majority of mutational processes (**Fig. 1D, Supplementary Fig. S1A-S1B**). While signature 1 and 6 are related to spontaneous deamination of 5-methylcytosine and DNA mismatch repair, respectively, which results in C > T transitions and predominantly occurs at NpCpG trinucleotides ^17,18^, other signatures specific to a study may be due to other endogenous mutational processes, treatment, or environment ^19^.

We also found that the recurrently mutated genes in our study were similar to TCGA and Li et al., and the overlap between these three studies is about 50% **(Supplementary Fig. S1C-1D, Supplementary Table S5-S6)**. Some cancer-related genes that have been reported in other populations (Hong Kong and Russian) were also found frequently mutated in our cohorts, including *TP53, LRP1B, PTPRT, ARID1A, FAT4, FAT1, APC, SMAD4, CDKN2A, CDH1, PIK3CA, RHOA*, and *PTEN*^7,8,15,20^

An analysis of copy number alterations of these 74 samples showed that most chromosome arms had undergone copy number gain or loss, with frequent amplified regions including 1q, 6p, 7, 8q, 13q, 20 (frequencies from 12% to 64%), and frequent losses observed on chromosomes 4, 14q, 18q, 19, 21q, 22q (frequencies from 16% to 43%) (**Supplementary Fig. S2A**). These overall somatic copy number variant (sCNV) patterns are consistent with previously published studies on GC ^7,8,20,21^. We identified 156 focal amplifications and 69 focal deletions, in well-known oncogenes, such as *ERBB2, CCNE1, KRAS, MYC, EGFR*, and *CDK6*, and cancer-related genes such as *GATA4, GATA6, CD44* and *ZNF217* (**Supplementary Table S7-S8**). Some tumor suppressor genes were identified in focal deleted regions, such as *CDKN2A, FAT1* and *SMAD4* (**Supplementary Fig. S2B**). These results are consistent with other studies such as TCGA and Wang et al. ^7,8^. Overall, we found 155 cancer genes amplified or deleted in our samples, in which half of them (78 genes) have been reported by TCGA or Wang et al. (**Supplementary Fig. S2C, Supplementary Table S9**), the other half (77 cancer genes) with sCNVs identified in our study could be further confirmed for their involvement in the development of gastric cancers.

### Mutation Load (TMB and TNB) of Chinese GC patients

TMB and specific neo-antigens have been reported as genomic biomarkers with the potential to impact cancer immunotherapy ^22^. Therefore, we next investigated the TMB and TNB of GC and the association with mutations in 17 DNA repair genes, such as *POLE, BRCA2, MSH3*. Across the entire GC dataset, the median TMB was 2.99 mutations/Mb, with a range of 0-50.57 mutations/Mb. Out of 74 samples, 8 (10.81%) had a high TMB (TMB > 10 mutations/Mb), 66 (89.19%) a low TMB (TMB <= 10 mutations/Mb). 61 (82.43% of 74) samples were successfully predicted neoantigens and 13 samples were failed due to the failure of HLA prediction in these samples (**Supplementary Table S10**). The median TNB was 2.47 neoantigens/Mb, with a range of 0.03-12.17 neoantigens/Mb. TNB is strongly associated with TMB (Pearson’s test, p= 3.631e-12, correlation = 0.75, **Fig. 2A**) and Missense Mutation Burden (MMB, Pearson’s chi-squared test, p = 2.242e-13, correlation = 0.78, **Fig. 2B**).

**Figure 2.**
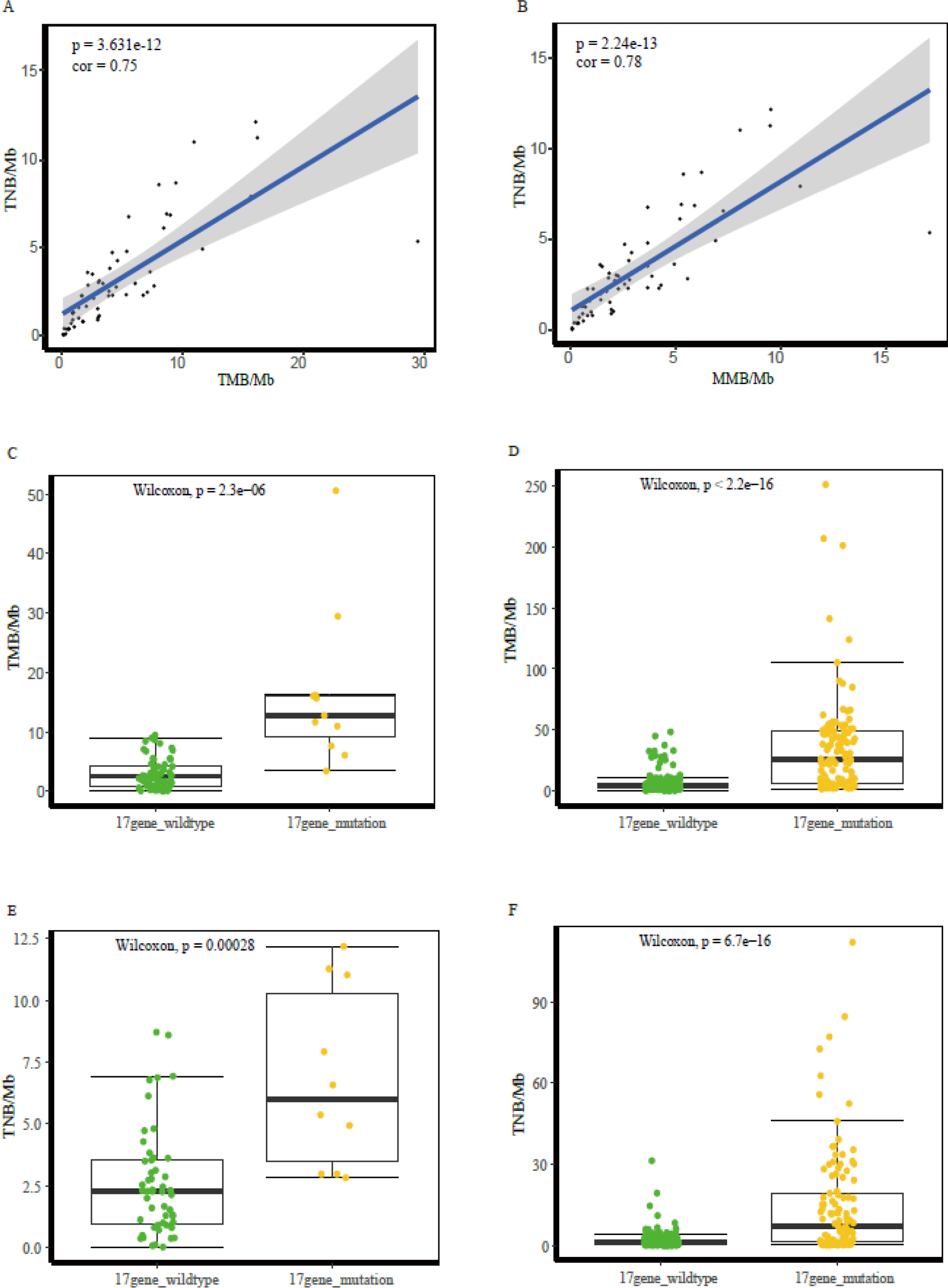
Association of mutations in 17 DNA repair genes with TMB and TNB. **(A)** The correlation between TMB and TNB. **(B)** The correlation between MMB and TNB. **(C-D)** The TMB comparison between wildtype and mutated samples associated with 17 DNA repair genes in this study(C) and the TCGA cohorts(D). **(E-F)** The TNB comparison between wildtype and mutated samples associated with 17 DNA repair genes in this study(E) and the TCGA cohorts(F).

In our study, samples with mutations in five DNA repair genes (5/17), including *CHEK1, MSH3, POLE, POLQ*, and *RAD50*, had a significantly higher TMB than wildtype samples, while samples with mutations in all these seventeen DNA repair genes (17/17) had a higher TMB than wildtype samples in TCGA dataset (**Table 2**). If we considered these 17 genes as a panel, we found that all samples with mutations of any of the 17 DNA repair genes, had significantly higher TMB than wildtype samples, both in our dataset (Wilcoxon test, p= 2.3e-6) and the TGCA dataset (Wilcoxon test, p < 2.2e-16) (**Fig. 2C-2D**).

**Table 2.** The association of TMB and genomic alterations in DNA repair genes

| Gene | This study |  |  | TCGA |  |  |
| --- | --- | --- | --- | --- | --- | --- |
|  | Mutated | Wildtype | P | Mutated | Wildtype | P value |
|  | Median(range), mutations/Mb | Median(range), mutations/Mb | value | Median(range), mutations/Mb | Median(range), mutations/Mb |  |
| BRCA2 | 26.99(3.40-50.57) | 2.97(0.00-29.43) | 0.17 | 43.94(1.41-251.25) | 4.045(0.03-105.19) | 3.00E-14 |
| CHEK1 | 20.18(10.93-29.43) | 2.97(0.00-50.57) | 0.032 | 127.39(5.13-251.25) | 4.31(0.03-141.16) | 0.00097 |
| CLK2 | 16.17(16.17-16.17) | 2.97(0.00-50.57) | 0.11 | 46.84(3.16-105.19) | 4.28(0.03-251.25) | 2.70E-05 |
| DCLRE1B | 15.63(15.63-15.63) | 2.97(0.00-50.57) | 0.13 | 141.16(6.94-251.25) | 4.295(0.03-123.94) | 4.60E-05 |
| DDB1 | 17.75(6.07-29.43) | 2.97(0.00-50.57) | 0.075 | 41.08(2.03-105.19) | 4.22(0.03-251.25) | 1.60E-05 |
| ERCC4 | 29.43(29.43-29.43) | 2.97(0.00-50.57) | 0.1 | 62.13(3.94-251.25) | 4.28(0.03-141.16) | 1.20E-05 |
| FANCI | 16.03(16.03-16.03) | 2.97(0.00-50.57) | 0.12 | 53.955(2.75-251.25) | 4.31(0.03-206.97) | 0.00068 |
| MSH3 | 40(29.43-50.57) | 2.97(0.00-16.17) | 0.017 | 56.345(2.59-201.19) | 4.28(0.03-251.25) | 0.00023 |
| PMS1 | 29.43(29.43-29.43) | 2.97(0.00-50.57) | 0.1 | 37.56(1.94-123.94) | 4.31(0.03-251.25) | 0.0059 |
| POLE | 15.9(15.63-16.17) | 2.97(0.00-50.57) | 0.027 | 49.75(2.97-251.25) | 4.075(0.03-123.94) | 6.10E-14 |
| POLN | 12.77(12.77-12.77) | 2.97(0.00-50.57) | 0.15 | 52.19(1.97-206.97) | 4.28(0.03-251.25) | 6.60E-05 |
| POLQ | 29.1(7.63-50.57) | 2.97(0.00-29.43) | 0.047 | 43.94(4.00-251.25) | 4.06(0.03-105.19) | 3.10E-13 |
| RAD50 | 33.37(16.17-50.57) | 2.97(0.00-29.43) | 0.019 | 39.685(2.22-251.25) | 4.31(0.03-206.97) | 0.0022 |
| RBBP8 | 29.43(29.43-29.43) | 2.97(0.00-50.57) | 0.1 | 30.66(22.41-42.28) | 4.295(0.03-251.25) | 0.001 |
| RRM2B | 11.63(11.63-11.63) | 2.97(0.00-50.57) | 0.16 | 33.63(11.47-46.84) | 4.345(0.03-251.25) | 0.031 |
| SHPRH | 12.77(12.77-12.77) | 2.97(0.00-50.57) | 0.15 | 49.685(2.66-251.25) | 4.19(0.03-206.97) | 1.90E-08 |
| TP53BP1 | 15.63(15.63-15.63) | 2.97(0.00-50.57) | 0.13 | 31.39(2.03-206.97) | 4.19(0.03-251.25) | 2.40E-05 |

We found no significant difference in TNB between wildtype samples and mutated samples of DNA repair genes (16/17) except for *POLE* (p = 0.03) (**Table 3)**. However, samples with mutations in DNA repair genes (13/17), had significantly higher TNB than wildtype samples in TCGA dataset. Similarly, if we considered these 17 genes as a panel, all samples with mutations of any of the 17 DNA repair genes, had significantly higher TNB than wildtype samples, both in our dataset (Wilcoxon test, p = 0.00028) and the TGCA dataset (Wilcoxon test, p = 6.7e-16) (**Fig. 2E-2F**). Interestingly, we found that the samples with mutations of any of the 17 DNA repair genes, had a shorter disease-free survival (DFS) than wildtype samples in our dataset (**Supplementary Fig. S3A**). These results indicate that mutations of DNA repair genes can affect TMB and TNB, and lead to a poor prognosis.

**Table 3.** The association of TNB and genomic alterations in DNA repair genes

| Gene | This study |  |  | TCGA |  |  |
| --- | --- | --- | --- | --- | --- | --- |
|  | Mutated | Wildtype | P value | Mutated | Wildtype | P value |
|  | Median(range), mutations/Mb | Median(range), mutations/Mb |  | Median(range), mutations/Mb | Median(range), mutations/Mb |  |
| BRCA2 | 2.97(2.97-2.97) | 2.4(0.03-12.17) | 0.73 | 12.67(0.06-111.91) | 0.94(0.03-72.50) | 9.5E-09 |
| CHEK1 | 8.2(5.37-11.03) | 2.33(0.03-12.17) | 0.06 | 21.72(0.31-111.91) | 1(0.03-84.47) | 0.1 |
| CLK2 | 11.27(11.27-11.27) | 2.4(0.03-12.17) | 0.11 | 14.28(1.53-72.50) | 1(0.03-111.91) | 0.77 |
| DCLRE1B | 7.93(7.93-7.93) | 2.4(0.03-12.17) | 0.16 | 39.09(0.12-111.91) | 1.00(0.03-84.47) | 0.0065 |
| DDB1 | 4.17(2.97-5.37) | 2.33(0.03-12.17) | 0.31 | 17.28(0.03-72.50) | 1.00(0.03-111.91) | 0.0018 |
| ERCC4 | 5.37(5.37-5.37) | 2.4(0.03-12.17) | 0.29 | 8.31(0.12-111.91) | 1.00(0.03-77.03) | 0.02 |
| FANCI | 12.17(12.17-12.17) | 2.4(0.03-11.27) | 0.094 | 7.565(1.38-52.34) | 1(0.03-111.91) | 0.014 |
| MSH3 | 5.37(5.37-5.37) | 2.4(0.03-12.17) | 0.29 | 16.31(0.41-77.03) | 1.00(0.03-111.91) | 0.0039 |
| PMS1 | 5.37(5.37-5.37) | 2.4(0.03-12.17) | 0.29 | 12.09(0.38-55.66) | 1(0.03-111.91) | 0.032 |
| POLE | 9.6(7.93-11.27) | 2.33(0.03-12.17) | 0.03 | 9.97(0.06-111.91) | 0.94(0.03-84.47) | 7.5E-07 |
| POLN | 6.57(6.57-6.57) | 2.4(0.03-12.17) | 0.24 | 14.205(0.03-111.91) | 1(0.03-84.47) | 0.072 |
| POLQ | 2.83(2.83-2.83) | 2.4(0.03-12.17) | 0.84 | 15.16(0.34-111.91) | 0.94(0.03-72.50) | 4.6E-10 |
| RAD50 | 11.27(11.27-11.27) | 2.4(0.03-12.17) | 0.11 | 18.875(0.50-52.34) | 1(0.03-111.91) | 0.0029 |
| RBBP8 | 5.37(5.37-5.37) | 2.4(0.03-12.17) | 0.29 | 12.81(0.31-45.84) | 1.00(0.03-111.91) | 0.014 |
| RRM2B | 4.93(4.93-4.93) | 2.4(0.03-12.17) | 0.32 | 13.50(1.16-28.53) | 1.00(0.03-111.91) | 0.085 |
| SHPRH | 6.57(6.57-6.57) | 2.4(0.03-12.17) | 0.24 | 15.09(0.31-52.34) | 0.985(0.03-111.91) | 5.3E-05 |
| TP53BP1 | 7.93(7.93-7.93) | 2.4(0.03-12.17) | 0.16 | 10.545(0.16-111.91) | 1(0.03-72.50) | 8.1E-04 |

Patients that were older (age >= 60) have significantly higher TMB (p = 0.0021) and TNB (p = 0.034) than younger patients (age < 60) (**Supplementary Fig. S3B-3C**). and male patients tend to carry more mutations than female patients (p = 0.034), but the difference in TNB was not significant (p = 0.82) (**Supplementary Fig. S3D-3E**).

### Genomic alterations associated with PM

The patients with PM had a worse prognosis than those without PM (p = 0.0034, **Fig. 3A**). To determine if there are genes specifically associated with PM, we identified 18 genes (Fisher exact test, p < 0.05) which showed moderate enrichment in the 22 patients who developed PM after surgery **(Fig. 3B)**. Using the KOBAS online database (http://kobas.cbi.pku.edu.cn), we found that these genes are enriched in cell adhesion by Gene Ontology (corrected p < 0.05). We found that all samples with mutations in any of 18 PM associated genes, had significantly higher TMB and TNB than wildtype samples, both in this study (**Fig. 3C-3D**) and the TCGA dataset (**Supplementary Fig. S4A-4B**). All samples with mutations of any 18 PM associated genes, had shorter DFS than wildtype samples in this study (**Supplementary Fig. S4C**). Furthermore, we found that the amplification of several regions is enriched in PM patients, and two of them (1p36.21 and Xq26.3) are associated with a worse prognosis (**Fig. 3D-3E**). Interestingly, the 1p36.21 region contains a gene family named *PRAME* (preferentially expressed antigen of melanoma), which is expressed in many cancers and was functions in reproductive tissues during development ^23^.

**Figure 3.**
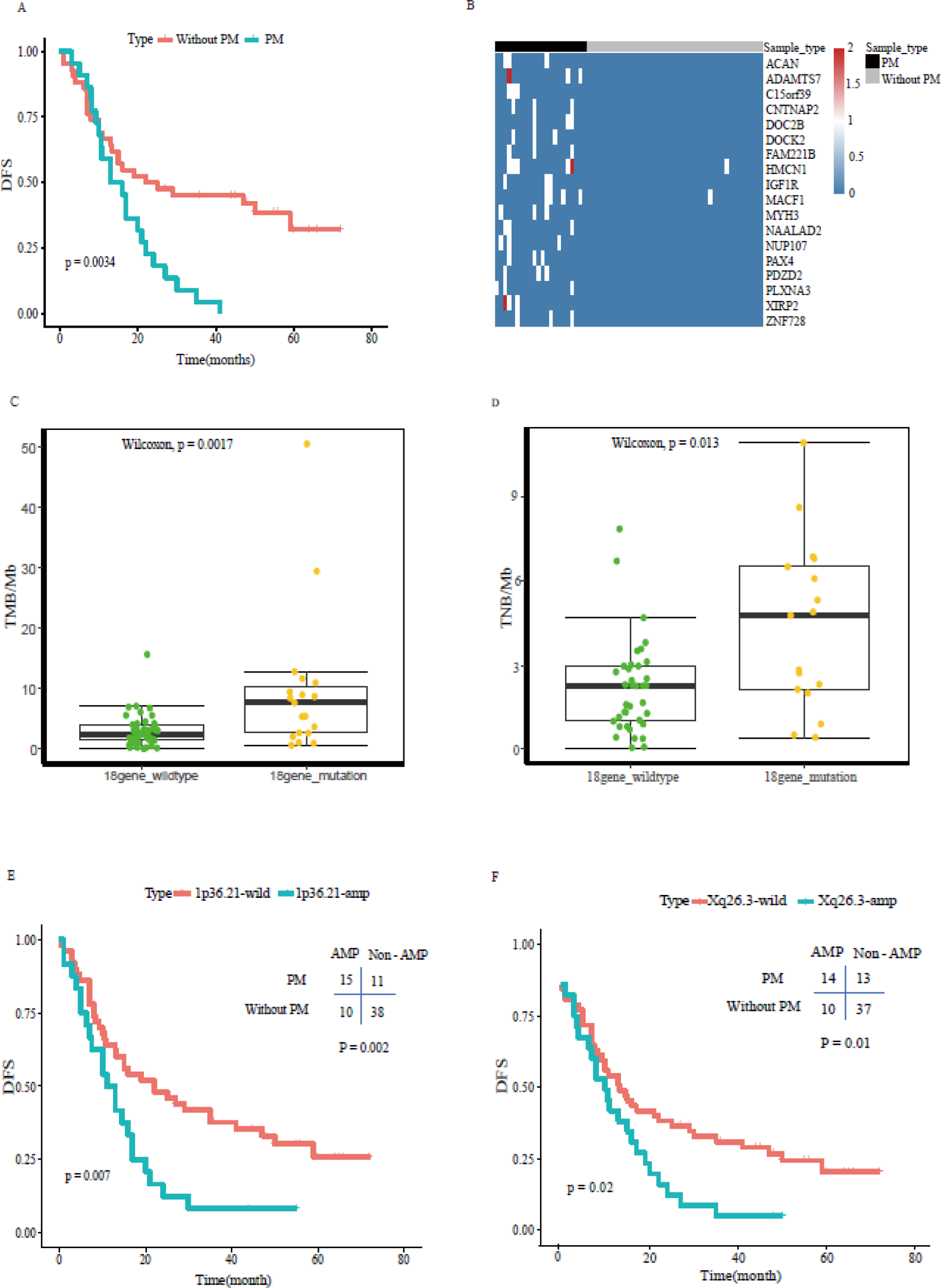
Genomic alterations associated with PM. **(A)** Kaplan-Meier plots for DFS in PM and not-PM patients. **(B)** Genes which enriched in PM patients. Fisher exact test, P<0.05. **(C-D)** The TMB and TNB comparison between wildtype and mutated samples of 18 PM associated genes in this study. **(E-F)** Kaplan-Meier plots for DFS in 1p36.21 and Xq26.3 for wildtype and mutated patients.

## Discussion

To our knowledge, this study is the first to assess both TMB and TNB in Chinese patients with GC and to describe the mutational profiles associated with increased TMB and TNB. It is believed that tumors with high TMB are more likely to harbor neoantigens that makes them vulnerable to attack by the immune system ^24^. In our study, we also found that tumors with high TMB tends to harbor high TNB, and the correlativity between TMB and TNB is significant. Mutations in 17 DNA repair genes are associated with high TMB and TNB both in our study and the TCGA cohorts, and the statistical difference of TMB between the mutated and wildtype group is significant in TCGA GC samples. Interestingly, we found that patients harboring mutations of these genes tend to have shorter DFS (**Supplementary Fig. S3A**), and these patients may benefit from alternative therapies.

Age is a major risk for cancer, and mutations accumulate with age ^7^. In our study, we found that older GC patients (age >= 60) harbor more mutations and neoantigens than younger patients (p = 0.0021 and p = 0.034, respectively). Interestingly, male patients also carry more mutations than female patients, which represent different risk factors.

PM can lead to bowel obstruction or malignant ascites, resulting in a poor prognosis and decline in the quality of life, so it is important to identify risk factors for PM ^4^. We found mutations in 18 genes are associated with PM, suggesting that these genes are potential markers of PM. We also found 2 regions, 1p36.21 and Xq26.3, that are amplified in PM patients, and associated with a poorer DFS. Due to the limited sample size of our study, further studies should be conducted to confirm this assocaition.

## Materials and Methods

### Patient cohort

This study was approved by the Ethical Committee of the Union Medical College Hospital Affiliated of Fujian Medical University and carried out according to the approved guidelines. In total, 300 cases with sufficient clinical pathological information were provided;155 of which with pathological paraffin blocks were selected for WES (**Supplementary Table S1**). Samples of cancer and adjacent normal tissues were taken from each case at the same time, a total of 6 FFPE sections with size of 10 μm in 1 cm × 1 cm and tumor content of more than 50% were selected. Of the selected samples, 74 were successful for subsequent library construction and sequencing.

### WES library construction and next-generation sequencing

The genomic DNA of FFPE samples was randomly fragmented and the size of the library fragments was mainly distributed between 150bp and 250bp. The end repair of DNA fragments was performed, and an “A” base was added at the 3’-end of each strand. Adapters were then ligated to both ends of the end repaired dA tailed DNA fragments for amplification and sequencing. Size-selected DNA fragments were amplified by ligation-mediated PCR, purified, and whole-exome capture was performed using the BGI Human All Exon V4 kit. Captured products were then circularized. Rolling circle amplification (RCA) was performed to produce DNA Nanoballs (DNBs). Each resulting qualified captured library was then loaded on BGISEQ-500 platform and pair-end 50bp or pair-end 100bp sequencing was conducted for each captured library. We sequenced an average of 1,533,107,107 reads for each sample, after reads quality filtering and duplication removing, the sequencing depths for FFPE tumors and corresponding normal tissues were 117× and 92× on averages, respectively (**Supplementary Table S11-S14**) **Identification of somatic mutations.** The sequencing data processing and variants detection pipeline is shown in **Supplementary Fig. S5.** Reads containing sequencing adapters and low-quality reads were removed using SOAPnuke software ^25^. Then the high-quality data of each sample was mapped to the human HG19 reference genome and the duplicate reads were removed with Edico software (http://edicogenome.com/dragen-bioit-platform/). To ensure accurate variant calling, local realignment around Indels and base quality score recalibration was performed using GATK ^26,27^. Then the sequencing depth and coverage for each sample were calculated based on the alignments, and samples with low coverage or depth were re-sequenced on the same library to achieve enough sequencing depth.

SSNVs and sIndels were detected using the MuTect ^28^ and Varscan2 software ^29^, respectively. Then these mutations (sSNVs and sIndels) were annotated with ANNOVAR ^30^ and followed by several filtering steps to remove potential false positives and obtain reliable results. For MuTect, in addition to the build-in filters, the following filtering criteria were applied: (1) total read count in tumor and normal DNA >= 10; (2) mutation allele fraction >= 10% and >= 5 reads that support this mutation; (3) mutation site is at least five bases away from the end of the read; (4) the SNV was not encompassed in short repeat regions; (5) presence of variant on both strands and the distribution of reads supporting this variant on the two strand is not biased; (6) the frequency of variant is less than 0.5% at 1,000 Genomes (1000G) database (http://www.1000genomes.org), Exome Sequencing Project (ESP) 6500 database (http://evs.gs.washington.edu/EVS) or Exome Aggregation Consortium (ExAC) database (http://exac.broadinstitute.org). For Varscan2, in addition to the built-in filters, the following filtering criteria were applied: (1) coverage >= 10 in normal DNA and coverage >= 10 in tumor DNA; (2) variant frequency >= 15%; (3) the Indel was not encompassed in short repeat regions; (4) the frequency of Indel is less than 0.5% at 1,000 Genomes (1000G) database, Exome Sequencing Project (ESP) 6500 database and Exome Aggregation Consortium (ExAC) database. The final mutation results were list in **Supplementary Table S2.**

SCNVs were detected by the CNV workflow tools within GATK4 (https://github.com/broadinstitute/gatk). The FFPE normal samples were used as control to identify tumor-specific genomic alterations. Then the copy-number segment data was used as input to the GISTIC2 program ^31^ to detect recurrently amplified or deleted genomic regions. GSITIC2 analysis was performed using the default parameters.

### Confirmation of mutations

36 mutation sites, containing 21 cancer gene mutations and 15 mutations in PM samples specific genes were randomly selected for mass spectrometry validation. In total, 34 mutations were validated by the MassARRAY platform (including mutations that not been detected before, such as mutations in *NUP107*), with a 94% validation rate. We considered validation a success when both the tumor and normal genotype generated by MassARRAY platform were the same as the sequencing result, and failure if the genotype called by mass spectrometry was not the same as sequencing.

### Neoantigen prediction

SSNV mutations were used to predict neoantigens by NetMHC, NetMHCpan, PickPocket, PSSMHCpan and SMM ^32^. The poor-quality peptides were removed according to two criteria: (1) IC50 < 500 in at least in three tools; (2) MT score < WT score for each peptide.

### Statistical methods

A Wilcoxon test was used to analyze the significance of the association of TMB and TNB with DNA repair genes, PM associated genes, patient age and patient gender. The Fisher exact test was used to analyze the significance of associations of the number of gene mutations with PM and not-PM. All tests were two-sided, and statistical significance was set at p < 0.05. The analysis of the correlation between mutation burden and neoantigen burden was made by Pearson’s chi-squared test. All statistical analyses were performed with RStudio software (Version 3.5.1)

### Data availability statement

The data reported in this study are available in the CNGB Nucleotide Sequence Archive (CNSA: https://db.cngb.org/cnsa;accession number CNP0000159).

## Acknowledgements

We thank the department of pathology at Union Medical College Hospital Affiliated to Fujian Medical University for their assistance in sample and data collection. We also thank Lei Chen and M. Dean for their constructive advices on the manuscript. We would like to thank Yun Zhao, Xuehui Tang and Lei Ge for their administrative support. This work was supported by the Critical Patented Project of The Science & Technology Bureau of Fujian Province, China (grant number 2013YZ0002-2), the Joint Project of the Natural Science and Health Foundation of Fujian Province, China (grant number 2015J01397) and the Shenzhen Science and Technology Program (JCYJ20170817145845968).

## Author Contributions

Q.C., X.-Q.Z. and C.C. conceived the experiments. X.-Q.Z., C.-M.S., Q.-L.L. and Y.Z. conducted the experiments. C.C., X.-C.H., Z.-Y.Z., Z.-K.Z., J.-W.Z., B.L. S.-Y.L. and S.Q. analyzed the results. F.W., R.-P.W. and Z.-Q.H. provided patient specimens and conducted histopathological examinations. C.C. and X.-C.H. wrote the manuscript. All authors reviewed the manuscript.

## Additional Information

**Competing Interests:** The authors declare no competing interests.

